# Functional buffering via cell-specific gene expression promotes tissue homeostasis and cancer robustness

**DOI:** 10.1101/2021.07.26.453775

**Authors:** Hao-Kuen Lin, Jen-Hao Cheng, Chia-Chou Wu, Feng-Shu Hsieh, Carolyn Dunlap, Sheng-hong Chen

## Abstract

Functional buffering that ensures biological robustness is critical for maintaining tissue homeostasis, organismal survival, and evolution of novelty. However, the mechanism underlying functional buffering, particularly in multicellular organisms, remains largely elusive. Here, we developed an inference index (C-score) for Cell-specific Expression- BUffering (CEBU), whereby functional buffering is mediated via expression of buffering genes in specific cells and tissues in humans. By computing C-scores across 684 human cell lines using genome-wide CRISPR screens and transcriptomic RNA-seq, we report that C- score-identified putative buffering gene pairs are enriched for members of the same duplicated gene family, pathway, and protein complex. Furthermore, CEBU is especially prevalent in tissues of low regenerative capacity (e.g., bone and neuronal tissues) and is weakest in highly regenerative blood cells, linking functional buffering to tissue regeneration. Clinically, the buffering capacity enabled by CEBU can help predict patient survival for multiple cancers. Our results reveal CEBU as a buffering mechanism contributing to tissue homeostasis and cancer robustness in humans.

**Summary blurb:** We unveil a genome-wide functional buffering mechanism, termed Cell-specific Expression Buffering (CEBU), whereby gene expression contributes to functional buffering in specific cell types and tissues. We link CEBU to genetic interactions, tissue homeostasis and cancer robustness.

## Introduction

Robustness in biological systems is critical for organisms to carry out vital functions in the face of environmental challenges ^1, 2^. A fundamental requirement for achieving biological robustness is functional buffering, whereby the biological functions performed by one gene can also be attained via other buffering genes. Although functional buffering has long been regarded as a critical function contributing to biological robustness, the mechanisms underlying functional buffering remain largely unclear ^3^. Based on transcriptional regulation of buffering genes, functional buffering can be categorized as either needs-based buffering or intrinsic buffering. Needs-based buffering involves transcriptional activation of buffering genes only when the function of a buffered gene is compromised. To accomplish needs-based buffering, a control system must exist that senses compromised function and then activates expression of buffering genes. Needs-based buffering is often observed as genetic compensation in various biological systems including fungi, animals and plants ^3–6^. One classical needs-based buffering mechanism is genetic compensation among duplicated genes, whereby expression of a paralogous gene is upregulated when the function of the active duplicated gene is compromised ^7^. Genetic analyses of duplicated genes in *Saccharomyces cerevisiae* have revealed upregulation of gene expression in ∼ 10% of paralogs when cell growth is compromised due to deletions of their duplicated genes ^6, 8, 9^. Apart from duplicated genes, non-orthologous/analogous genes can also be activated for needs-based buffering ^10^.

For instance, inactivation of one growth signaling pathway can lead to activation of others for the coordination of cell growth and survival ^3, 7^. Such needs-based buffering genes have been documented as enabling unicellular/multicellular organisms to cope with environmental stresses ^3, 9^.

Recent genome-wide studies of duplicated genes in human cells have revealed another class of buffering mechanism whereby expression of buffering genes is not responsive to impaired function but is constitutively expressed, hereafter termed “intrinsic buffering” ^11–13^. In some duplicated gene families, the strength of paralogous gene expression determines the essentiality of their corresponding duplicated genes in human cell lines, i.e., the higher the expression of paralogous genes in a particular cell line, the less essential are their duplicated genes ^11–13^. This observation indicates that paralogs may buffer and contribute to the function of their duplicated genes in specific cells through their constitutive gene expression. In addition to duplicated gene families, gene essentiality can depend on inherent variability in the expression levels of other genes in the same pathway, suggesting that functionally analogous genes in the same pathway can also buffer each other ^14^. Despite these observations, it remains unclear what mechanism may give rise to this context-dependent constitutive expression of buffering genes and how such intrinsic buffering may function in multicellular organisms.

In this study, we directly investigated if cell- and tissue-specific gene expression can act as an intrinsic buffering mechanism (which we have termed “Cell-specific Expression-BUffering” or CEBU, **Fig. 1A**) to buffer functionally related genes in the genome, thereby strengthening cellular plasticity for cell- and tissue-specific tasks. To estimate buffering capability, we developed an inference index, the C-score, to identify putative gene pairs displaying CEBU. This index calculates the adjusted correlation between expression of a buffering gene and the essentiality of the buffered gene (**Fig. 1B**), utilizing transcriptomics data ^15^ and genome-wide dependency data from the DepMap project ^15, 16^ across 684 human cell lines. Our results suggest that CEBU-mediated intrinsic buffering plays a critical role in cell-specific survival, tissue homeostasis, and cancer robustness.

**Figure 1.**
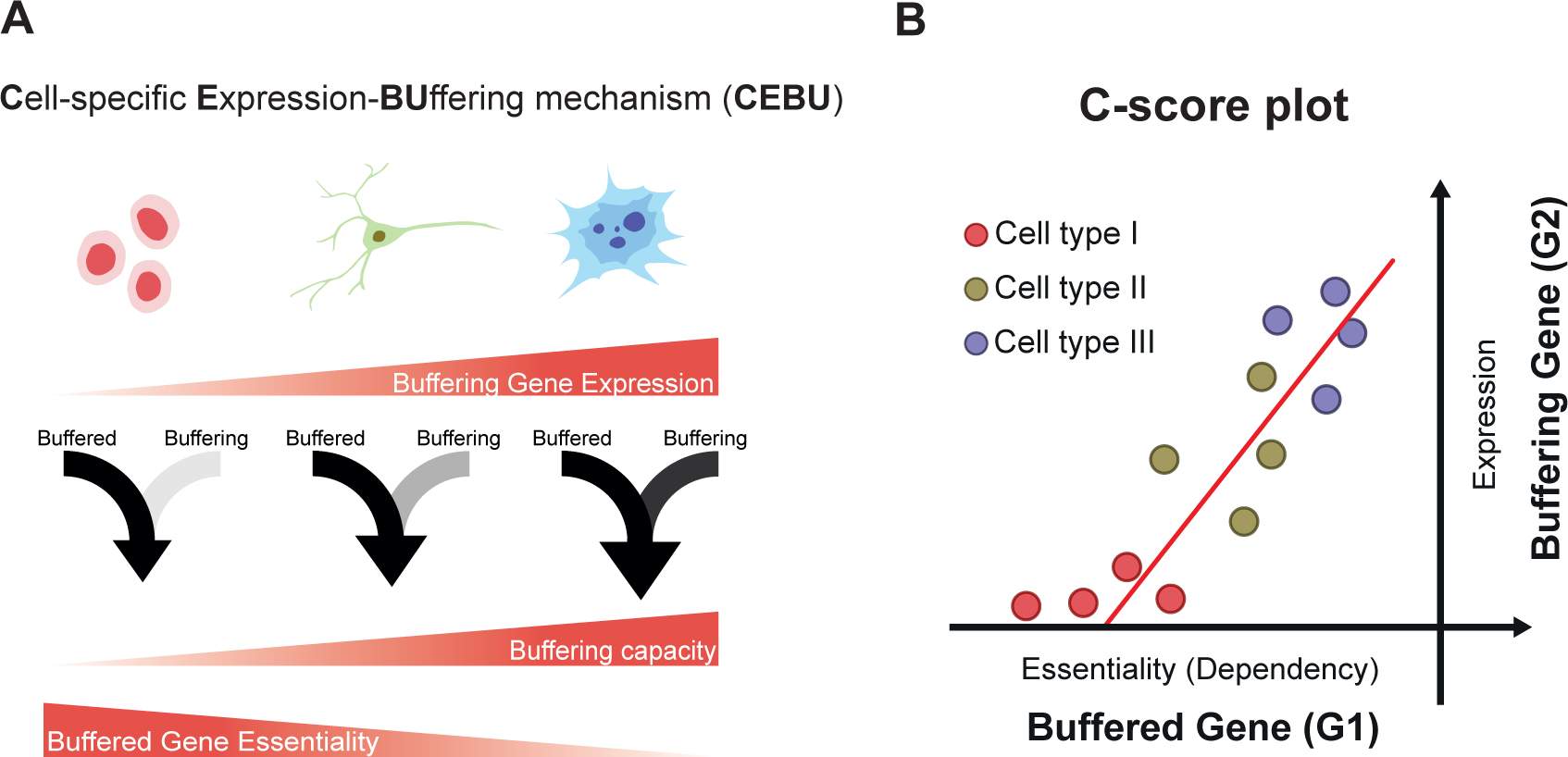
Genome-wide CEBU analysis using the C-score index. C-score plot: the x-axis is the dependency score of the buffered gene (G1) and the y-axis is the expression level of the buffering gene (G2). G1 is considered as being potentially buffered by G2, as quantified by C-score, which is an adjusted correlation for a given gene based on the C-score plot.

## Results

### Development of the C-score to infer cell-specific expression buffering (CEBU)

In seeking an index to infer intrinsic buffering operated via constitutive gene expression, we postulated a buffering relationship whereby the essentiality of a buffered gene (G1) increases when expression of its buffering gene (G2) decreases across different human cell lines (**Fig. 1**). Given that G2 expression differs among cell lines, the strength of buffering capacity varies across cell lines, thereby conferring on G1 cell-specific essentiality. This cell-specific expression buffering mechanism, here named CEBU, is the basis for our development of the C-score. The C-score of a gene pair is derived from the correlation between the essentiality of a buffered gene (G1) and expression of its buffering gene (G2) (see C-score plot, **Fig. 1B**), and is formulated as:

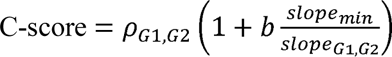

where *ρ* denotes the Pearson correlation coefficient between essentiality of G1 and expression of G2. Their regression slope (*slope_G1, G2_*) is normalized to *slope_min_*, which denotes the minimum slope of all considered gene pairs in the human genome (see **Methods**). The normalized slope can be weighted by cell- or tissue type-specific *b*. In our current analysis, *b* is set as 1 for a pan-cell- and pan-cancer-type analysis. Gene essentiality is represented by dependency scores (D.S.) from the DepMap project ^16^, where the effect of each gene on cell proliferation was quantified after its knockout using the CRISPR/Cas-9 approach. Specifically, a more negative D.S. reflects slower cell proliferation when the gene is knocked out, thus reflecting stronger essentiality. Expression data was obtained through RNA-seq ^15^. We anticipated that the higher the C-score of a gene pair, the more likely G2 would buffer G1 based on our proposed CEBU mechanism.

We conducted a genome-wide analysis to calculate C-scores for all gene pairs across 684 human cell lines. The calculated C-scores were compared to a bootstrapped null distribution generated by random shuffling of G2 expression among cell lines (**Fig. S1A**). The bootstrapped null distribution can be modeled as a normal distribution (**Fig. S1B**). For our analysis, we considered gene pairs to have a high C-score with a strong likelihood of intrinsic buffering when their C-scores were > 0.25 (0.058% of gene pairs in the human genome, significant with a *q*-value < 2.2e-16 after multiple testing correction, **Fig. S1A**). Based on our C-score definition, a high C-score should be indicative of marked variability in cell-specific essentiality and expression. Indeed, we observed higher variability in both G1 dependency and G2 expression for high C-score gene pairs (**Fig. S2A**). Nevertheless, high variation alone is insufficient to grant a high C-score. A high C-score requires consistent pairing between G1 and G2 across cell lines and, as anticipated, disrupting the pairing between G1 dependency and G2 expression by shuffling G2 expression amongst cell lines (without changing variability) abolished the C-score relationship (compare the right panel of **Fig. S2B** to the left one). Moreover, both mean G1 dependency and mean G2 expression in high C-score gene pairs were lower than those parameters in randomly selected gene pairs (**Fig. S2C**), implying that G1s and G2s in high C-score gene pairs tend to be more essential and less expressed, respectively.

### Characterization of C-score-inferred CEBU gene pairs

Since several duplicated gene pairs have been implicated as displaying functional buffering via gene expression ^11–13^, we characterized the duplicated genes among C-score-identified gene pairs. We found that duplicated gene pairs are enriched among gene pairs with C-scores > 0.255 (*p*-value = 0.05 using a hypergeometric test), suggesting that CEBU is a prevalent buffering mechanism among duplicated genes (**Fig. 2A**). Interestingly, the majority of high C-score gene pairs are non-duplicated (> 90%, **Fig. S3A**). In these cases, G2s may be functional analogs of the respective G1s, acting as surrogate genes. Accordingly, we examined if the high C-score gene pairs are more likely to participate in the same function or biological pathway or physically interact. To do so, we calculated the enrichment of curated gene sets in terms of Gene Ontology (GO) ^17^ and Kyoto Encyclopedia of Genes and Genomes (KEGG) ^18^ from the Molecular Signatures Database ^19^ (**Fig. 2B**). Gene pairs with high C-scores consistently exhibited greater functional enrichment. Likewise, we observed a monotonic increase in the enrichment of protein-protein interactions (PPI) [using the STRING ^20^ and CORUM ^21^ databases] between G1s and G2s in accordance with increasing C-score cutoff (**Fig. 2C**). The enriched functions include housekeeping functions such as regulating redox homeostasis, gene transcription, mRNA translation, as well as NTP synthesis (**Fig. 2D**). Moreover, some cancer-related pathways are also enriched in the C-score-identified buffering network, including the proto-oncogenes *EGFR* and *MYC* (**Fig. 2D**). Both duplicated and non-duplicated gene pairs contribute to the observed functional and PPI enrichments. However, notably, functional and PPI enrichment are primarily attributable to non-duplicated genes (compare **Fig. 2B and 2C** to **Fig. S3B and S3C**), indicating a strong likelihood for intrinsic buffering among analogous genes in the same pathway or proteins in the same protein complex. Thus, high C-score gene pairs are enriched in duplicated gene pairs, as well as non-duplicated gene pairs that are members of the same biological pathway and/or encode physically interacting proteins, supporting that CEBU (which is the basis of our C-score index) is the mechanism enabling intrinsic buffering between such gene pairs.

**Figure 2.**
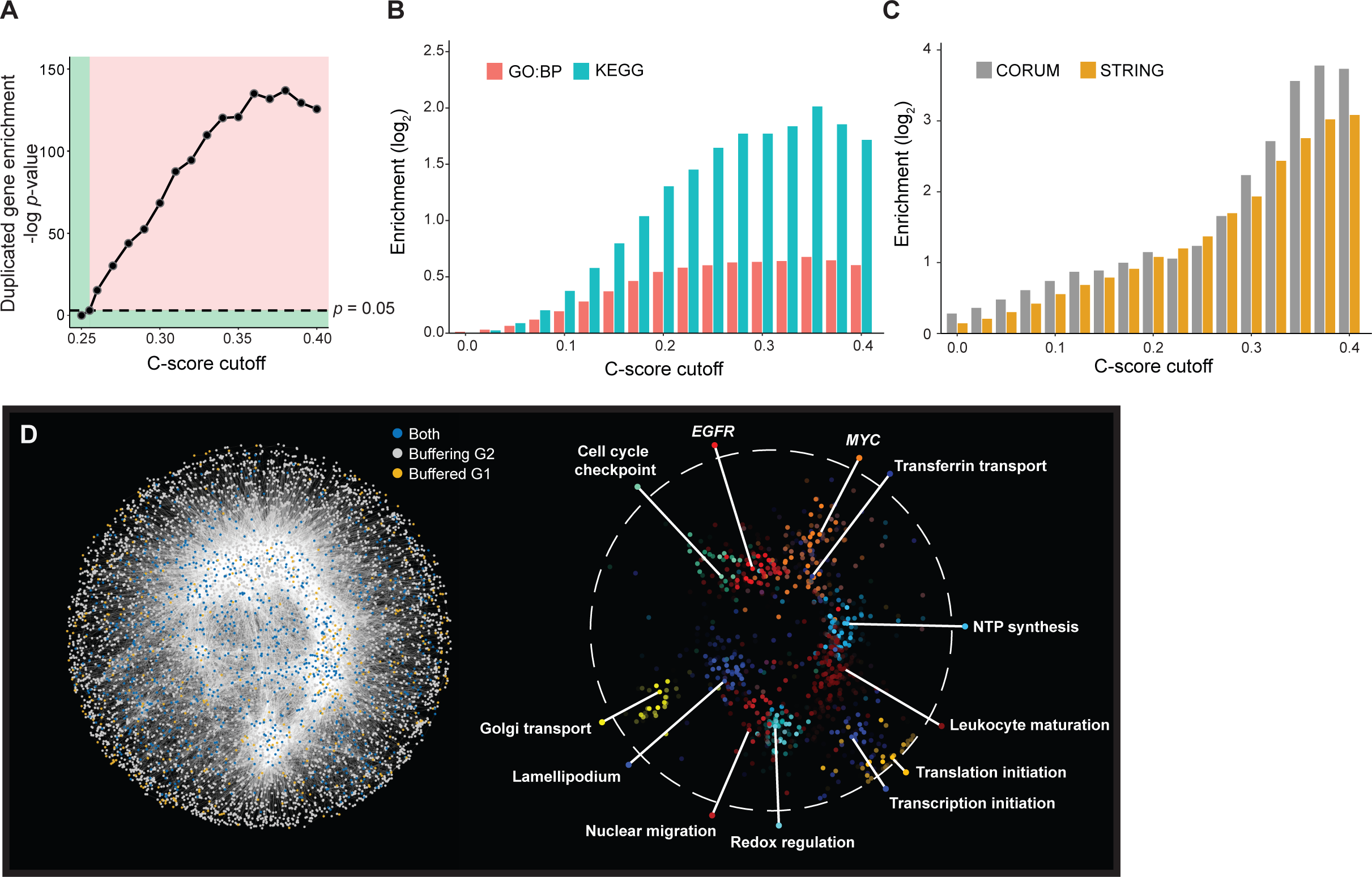
Properties of high C-score gene pairs. (A) Enrichment for duplicated genes as C-scores increase (hypergeometric test). The dashed line denotes the *p*-value of 0.05, where the corresponding C-score is 0.255. The red region (i.e., above the dashed line and equating to C-score > 0.255) indicates significant enrichment. The green region indicates lack of significance. (B-C) Functional enrichment of C-score gene pairs increases with C-score cutoff. (B) Enrichment of pairs of genes annotated with the same gene ontology biological process (GO:BP) term or KEGG pathway in C-score gene pairs. Enrichment increases with C-score cut-off. (C) Enrichment of pairs of genes with annotated protein-protein interactions from STRING and within the same protein complex from CORUM among C-score gene pairs. Enrichment increases with C-score cut-off. (D) Left: the buffering gene network is composed of 6,664 nodes and 42,754 edges with C-scores > 0.2536. Orange nodes represent buffered genes; grey nodes are buffering genes, and blue nodes are genes that are both buffered and buffering. Right: Clusters of GO-enriched biological functions in the buffering gene network.

### Experimental validation of C-score-inferred CEBU gene pairs

To validate putative C-score-inferred buffering gene pairs, we conducted experiments on the highest C-score gene pair, i.e., *FAM50A*-*FAM50B*, both members of which belong to the same duplicated gene family. Based on a C-score plot of *FAM50A-FAM50B* (**Fig. 3A**), we expected that *FAM50B* would display a stronger buffering effect on *FAM50A* for cell lines located at the top-right of the plot (e.g. A549 and MCF7) relative to those at the bottom-left (e.g. U2OS). Accordingly, growth of the cell lines at the top-right of the plot would be more sensitive to dual suppression of *FAM50A* and *FAM50B*. We used gene-specific small hairpin RNAs (shRNAs) to suppress expression of *FAM50A* and *FAM50B*, individually and in combination. Consistent with our expectations, we observed stronger growth suppression in the A549 and MCF7 cell lines relative to the U2OS cell line (**Fig. 3B**). Next, we quantified

**Figure 3.**
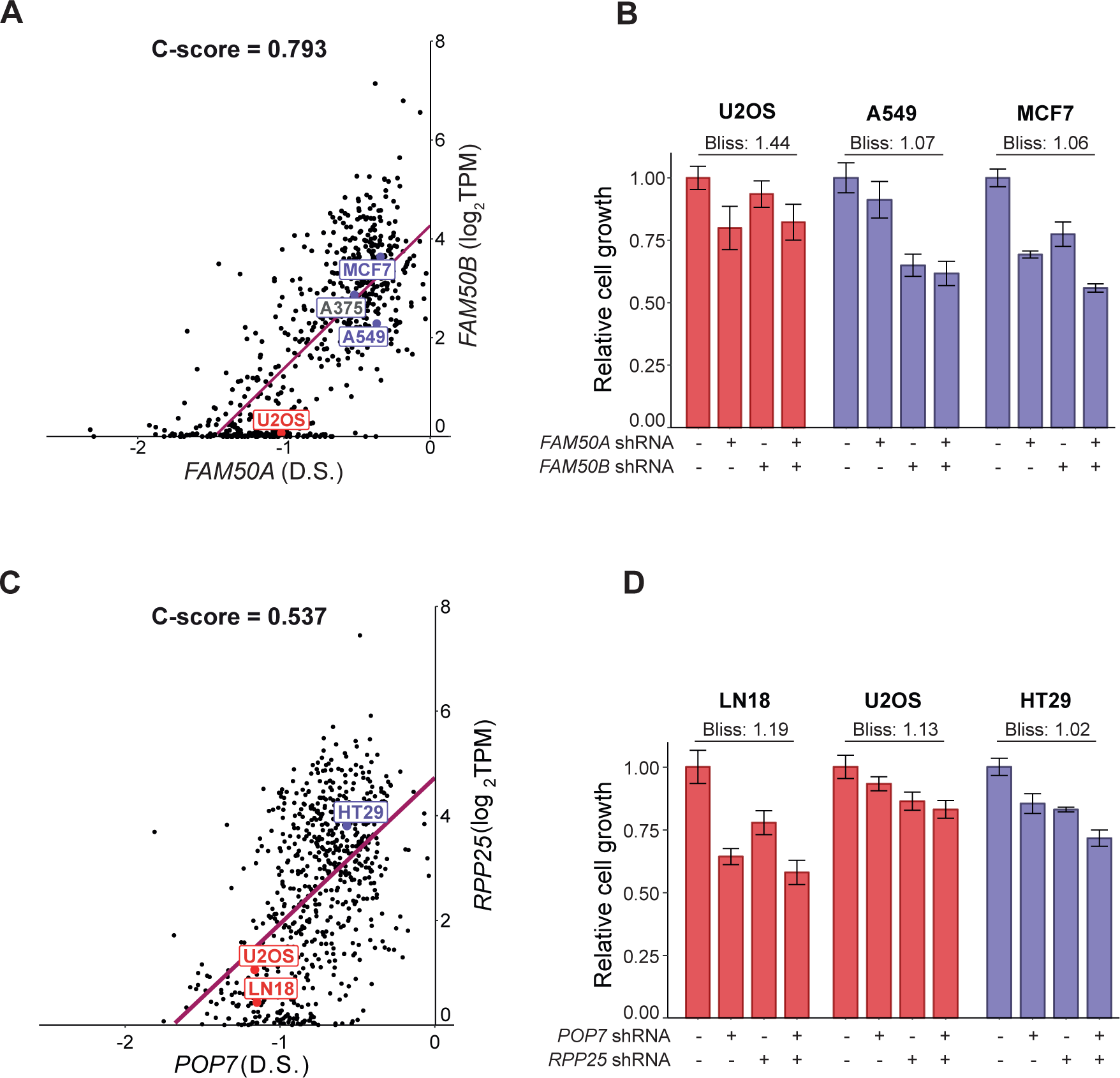
Experimental validation of cell-specific expression buffering between *FAM50A* and *FAM50B*, and *POP7* and *RPP25*. (A) C-score plot of the highest C-score gene pair, *FAM50A* (dependency score - D.S.) and *FAM50B* (G2) expression, labeled with the U2OS (predicted not synergistic), A549 (predicted synergistic), and MCF7 (predicted synergistic) cell lines. (B) Relative cell growth based on fold-change in confluency of the U2OS, A549, and MCF7 cell lines with or without *FAM50A* or *FAM50B* suppression. Bliss scores indicate strength of synergy between double suppression of *FAM50A* and *FAM50B* compared to either gene alone. Error bars indicate standard deviation of six technical repeats. (C) C-score plot of the non-duplicated gene pair, i.e., *POP7* dependency score and *RPP25* gene expression, labeled with the LN18 (predicted not synergistic), U2OS (predicted not synergistic), and HT29 (predicted synergistic) cell lines. (D) Relative cell growth based on fold-change in confluency of the HT29, U2OS and LN18 cell lines with or without shRNA-based *POP7* or *RPP25* suppression. Bliss scores indicate strength of synergy between double suppression of *POP7* and *RPP25* compared to either gene alone. Error bars indicate standard deviation of six technical repeats.

*FAM50A* and *FAM50B* genetic interactions in these three different cell lines by Bliss score ^22^, with lower scores indicating stronger synergistic interactions (see **Methods**). Indeed, the *FAM50A*-*FAM50B* gene pair in the A549 and MCF7 cell lines exhibited stronger synergy than in the U2OS cell line (Bliss score: 1.07 in A549, 1.06 in MCF7 and 1.44 in U2OS). Importantly, a recent study focusing on genetic interaction of duplicated genes identified the *FAM50A* and *FAM50B* gene pair as the most significant interacting duplicated gene pair in the human genome ^23^, further supporting the inference power of our C-score index. Moreover, the A375 cell line was used in that recent study, and it is predicted to display strong synergy based on our C-score plot of *FAM50A* and *FAM50B* (**Fig. 3A**).

Although duplicated genes are well recognized for their buffering relationship, there is limited evidence supporting intrinsic buffering among non-duplicated genes. Thus, we sought to experimentally examine a pair of non-duplicated genes with a high C-score, so we targeted the *POP7*-*RPP25* pair. These two genes encode protein subunits of the ribonuclease P/MRP complex. In the C-score plot of *POP7*-*RPP25* (**Fig. 3C**), the HT29 cell line lies in the top-right region and the U2OS and LN18 cell lines are in the bottom-left region, indicating a likelihood for a stronger buffering effect in the HT29 cell line. When we suppressed expression of *POP7* and *RPP25* using gene-specific shRNAs in these three cell lines, we observed that dual suppression of *POP7* and *RPP25* resulted in strong synergistic effects for the HT29 cell line but not for the U2OS or LN18 cell lines (Bliss scores for *POP7*-*RPP25* genetic interactions are 1.02 in HT29, 1.13 in U2OS, and 1.19 in LN18, **Fig. 3D**), indicating that C-score-inferred buffering gene pairs can be non-duplicated functional analogs in the same protein complex or duplicated genes of the same family.

### Tissue specificity of CEBU

One key feature of intrinsic buffering is cross-cell variation in the expression of buffering genes (G2s), which contributes to cell-specific dependency of the buffered genes (G1s) (**Fig. 1**). We hypothesized that the source of this cross-cell variation in G2 expression is embedded in the distinct transcriptional programs of different tissues. Therefore, we examined if the expression of high C-score G1s and G2s is tissue-specific. We calculated a tissue specificity index, *τ* ^24^, for each gene to establish if it displays low (low *τ*, broadly expressed across tissues) or high tissue specificity (high *τ*, only expressed in one or a few specific tissues). As shown in **Figure 4A**, G2s generally presented higher tissue specificity compared to G1s (significant with *t*-test, *p* < 2.2e-16) and compared to the control generated by randomly shuffling G2s across cell lines (**Fig. S4**). Together, these results indicate that G1s are generally expressed in the majority of cell types, whereas expression of G2s is more tissue- specific.

**Figure 4.**
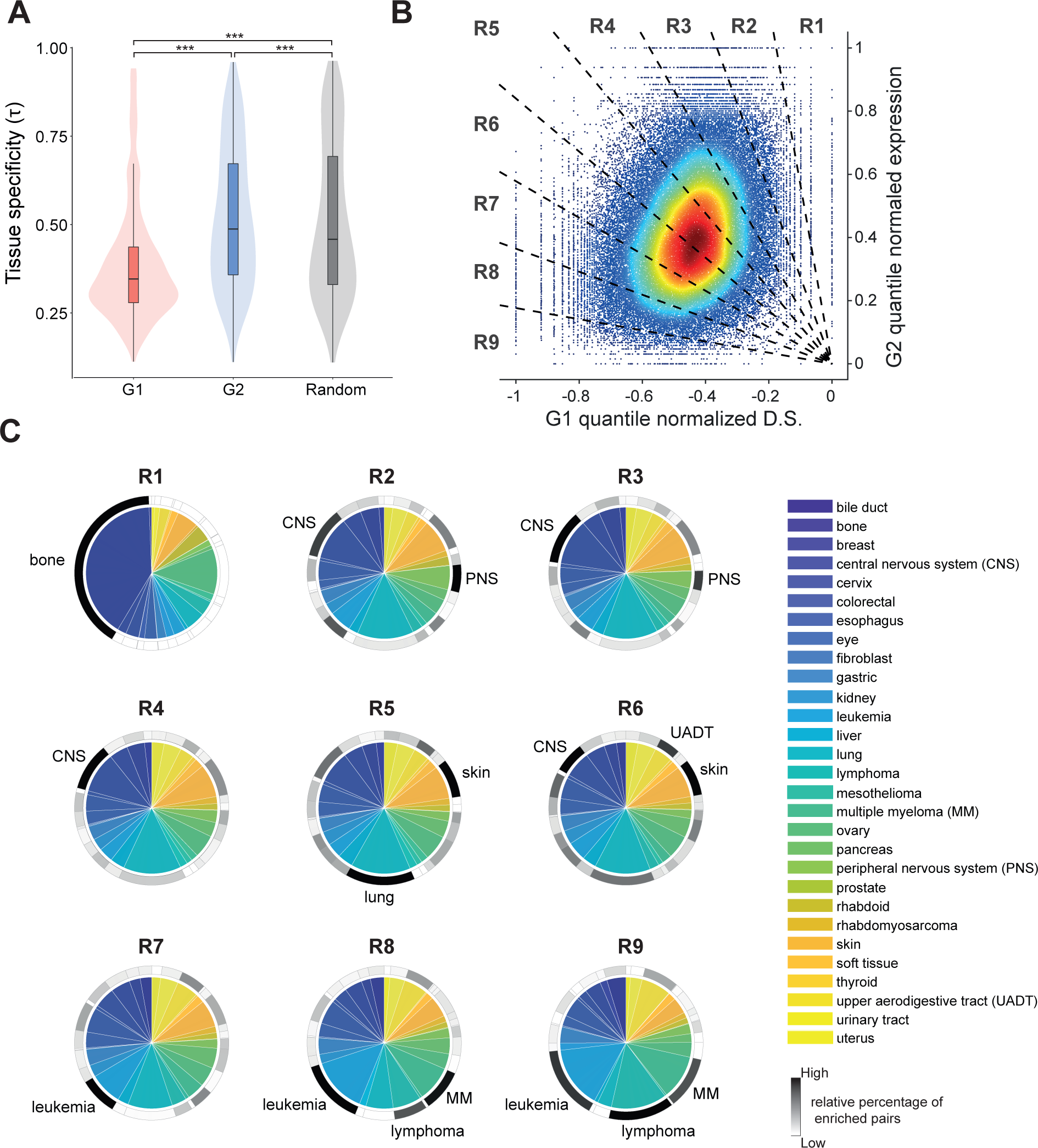
Tissue specificity of CEBU. (A) Tissue specificity (*τ* ) of G1 and G2 pairs. *τ* was calculated for G1 and G2 from high C- score gene pairs. Statistical significance was assessed by paired-*t* test. (B) The density plot of 100,000 randomly selected high C-score gene pairs. D.S. (G1s) and expression (G2s) were normalized to be between -1 and 0 or 0 and 1, respectively. The normalized C-score plot was divided equally into nine regions (R1-R9) by radiating lines out from zero. (C) Tissue/cell- type specificity of each region (R1-R9) of the normalized C-score plot. The colored pie charts indicate the proportion of each tissue/cell type in each of the regions. The greyscale rings around the pie charts represent the relative percentage of statistically enriched gene pairs for the corresponding tissue/cell types. The tissue/cell types with high percentages of enriched gene pairs (dark grey or black) are annotated for each region.

The pronounced tissue-specificity of G2 expression implies that CEBU acts as a type of tissue-specific intrinsic buffering. To further explore in which tissue types CEBU is more active, we generated normalized C-score plots for all high C-score gene pairs whereby the G1 dependency scores across all cell lines were quantile-normalized to be between -1 and 0 and the G2 expression values were normalized to be between 0 and 1 (**Fig. 4B**). We plotted these values against each other and then divided the resulting plot into nine equal regions by radiating lines out from zero (R1 to R9, **Fig. 4B**). As per the examples shown in **Fig. 3A, C**, tissue types displaying stronger CEBU-mediated buffering capacity would be enriched in the regions R1-R4, whereas those with low buffering capacity would predominate in regions R6- R9. Accordingly, considering a total of 29 tissue/cell types, we calculated the proportion of each tissue/cell type in each region of the plot in **Fig. 4B**, as well as the percentage of CEBU- enriched gene pairs for each tissue/cell type (see **Methods**). For each plot region, we observed that one to three tissue/cell types presented a high percentage of CEBU-enriched gene pairs (**Fig. 4C**). For example, for region R1, 98.0% of the CEBU-enriched gene pairs are highly expressed in cells derived from bone tissue (see grayscale ring surrounding the upper-left subplot of **Fig. 4C**), whereas region R9 encompasses a high percentage of strongly-expressing CEBU-enriched gene pairs in blood cells (lymphoma: 13.9%, leukemia: 10.6%, and multiple myeloma: 9.2%, see grayscale ring surrounding the bottom-right subplot of **Fig. 4C**). We also noted a few reoccurring tissue/cell types across regions of the plot reflecting high buffering capacity (central nervous system in R2, R3, and R4) or in low buffering regions (leukemia in R7, R8, and R9; lymphoma in R8 and R9) (**Fig. 4C**), indicating that particular tissue/cell types display a propensity for CEBU activity. These results support that CEBU reflects tissue-specific intrinsic buffering, and that whereas buffered G1s are generally expressed across tissue types, the buffering G2s are expressed in specific tissue/cell types, thereby contributing to tissue-specific functions.

### Harnessing C-score to calculate the buffering capacity of CEBU

As revealed by our experimental results in **Figure 3**, cell lines located in the upper right of a C-score plot are more sensitive to dual gene suppression, indicating a higher buffering capacity from G2s. To quantify G2 buffering capacities in various cells or tissues, we calculated buffering capacities as the relative G2 expression (compared to that of all other cell lines) of the cell line of interest adjusted by the C-score of the gene pair (**Fig. 5A** and **Methods**). We validated these C-score-derived buffering capacities as predictions of genetic interactions between G1 and G2 using experimental results from four independent studies in human cells (**Table S1)** ^25–28^. Using the receiver operating characteristic (ROC) curve to assess the performance of buffering capacity predictions, we observed that the resulting area under curve (AUC) is significantly larger than random (Mann-Whitney U test with *p*-value < 0.05, **Fig. 5B**). Furthermore, predictive performance increased for higher C-score cutoffs, as indicated by their increasing AUC (**Fig. 5B**). Moreover, the buffering capacity of CEBU is quantitatively correlated with the strength of genetic interaction. We observed a negative correlation between C-score-derived buffering capacities and experimentally validated genetic interactions (C-score cutoff = 0.25, correlation = -0.231, *p*-value = 0.034, **Fig. S5A**), and this correlation is stronger for higher C-score cutoffs (**Fig. S5B**). Even though this correlation coefficient of -0.231 is not strong (although it is statistically significant), the intrinsic variability associated with collating experimental results from four independent studies must be considered a contributory factor to weakening that correlation ^25–28^. Moreover, predictions of genetic interactions based on CEBU buffering capacity are robust even when different thresholds for calculating buffering capacity are applied (**Methods** and **Fig. S5C**). Accordingly, the CEBU mechanism can be used to infer genetic interactions in human cells. Since CEBU is reflective of tissue-specific intrinsic buffering (**Fig. 4**), we also quantified buffering capacity in various tissue/cell types. We calculated the average buffering capacity for each tissue/cell type based on high C-score gene pairs (**Fig. 5C**). In line with our enrichment analysis presented in **Figure 4**, the top three most buffered tissues are the central nervous system, bone and the peripheral nervous system. In contrast, blood cells—including multiple myeloma, lymphoma, and leukemia cell lines—exhibited the lowest buffering capacities.

**Figure 5.**
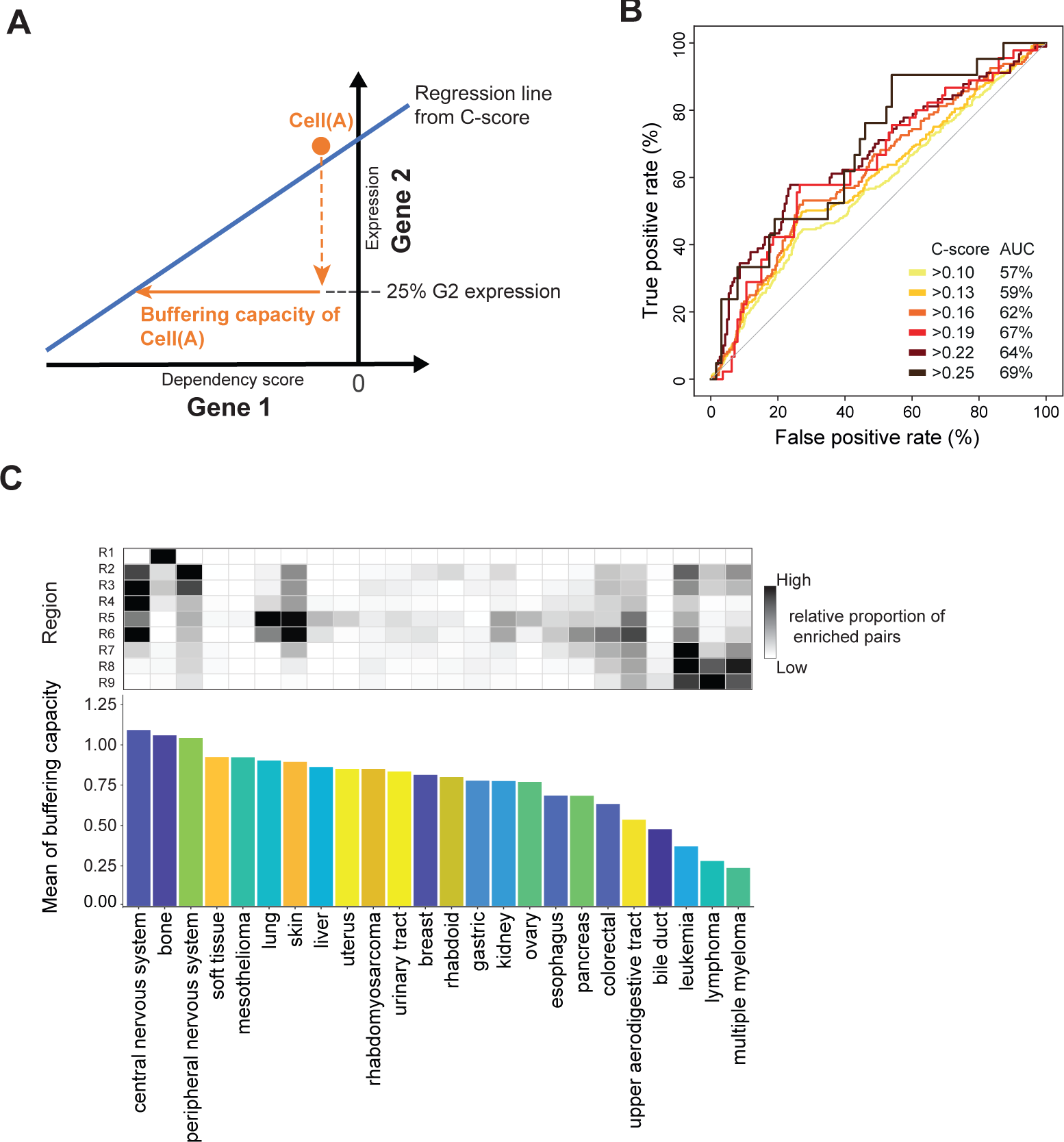
C-score-derived tissue-specific buffering capacity. (A) Illustration showing how cell-specific buffering capacities were derived from C-scores. Buffering capacity was calculated based on: 1) the regression line of the C-score for the gene pair; and 2) relative G2 expression (compared to that of all other cell lines) for the cell line of interest. See **Methods** for the formula for buffering capacity calculation. (B) Predictive performance shown as ROC curves for predicting genetic interactions using cell-specific buffering capacity. Prediction sets consist of 84 data-points (37 unique genetically interacting gene pairs) across 8 cell lines with a C-score cut-off of 0.25. (C) Mean buffering capacity for each tissue type (lower panel) and the corresponding proportion of enriched gene pairs for each region (upper panel).

### CEBU-mediated buffering capacity is indicative of cancer aggressiveness

Inspired by the proto-oncogenes we identified according to C-scores (**Fig. 2D**), we wondered if cancers in various tissues may take advantage of the buffering capacities endowed by the CEBU mechanism for robust proliferation. In other words, would higher CEBU-mediated buffering capacity render cancers more robust and aggressive, thereby resulting in a poorer prognosis? To test this hypothesis, we established a “ground-truth” of expression-based cancer patient prognosis by analyzing patient gene expression and survival data for all 30 available cancer types from The Cancer Genome Atlas (TCGA) ^29^. Here, we assessed differential patient survival against gene expression using Cox regression and controlling for clinical characteristics including age, sex, pathological stage, clinical stage, and tumor grade, followed by multiple testing correction (false discovery rate < 0.2). Then, we examined the performance of CEBU-mediated buffering capacity in terms of predicting the ground-truth dataset. As an example, in **Figure 6A** we present potential buffering to *NAMPT* of the NAD^+^ salvage pathway, where cancers may be addicted to this pathway ^30^. We discovered that the *NAMPT-CALD1* gene pair, comprising the *NAMPT* dependency score and *CALD1* gene expression, demonstrate a high C-score of 0.446, and its CEBU-mediated buffering capacity is high in CNS but low in blood cells. When we stratified patients based on *CALD1* expression, we observed a considerable difference in survival for patients suffering lower grade glioma (LGG – a cancer of the CNS, see **Table S2** for cross-referencing between cell lines and TCGA cancers and for the full names of cancer abbreviations), but not for patients with acute myeloid leukemia (LAML – a cancer of the blood, **Fig. 6B** left panel for LGG and right panel for LAML). Mean CEBU-mediated buffering capacity for the *NAMPT*:*CALD1* gene pair is 1.47 in the CNS (i.e. tissue/cell types displaying strong buffering capacity), but only -0.88 in leukemic blood cells (i.e. exhibiting weak buffering capacity) (**Fig. 6A**). Thus, based on our ground-truth dataset, the buffering capacity of the *NAMPT* and *CALD1* gene pair in different tissue/cell types can be used to predict patient survival for specific cancer types.

**Figure 6.**
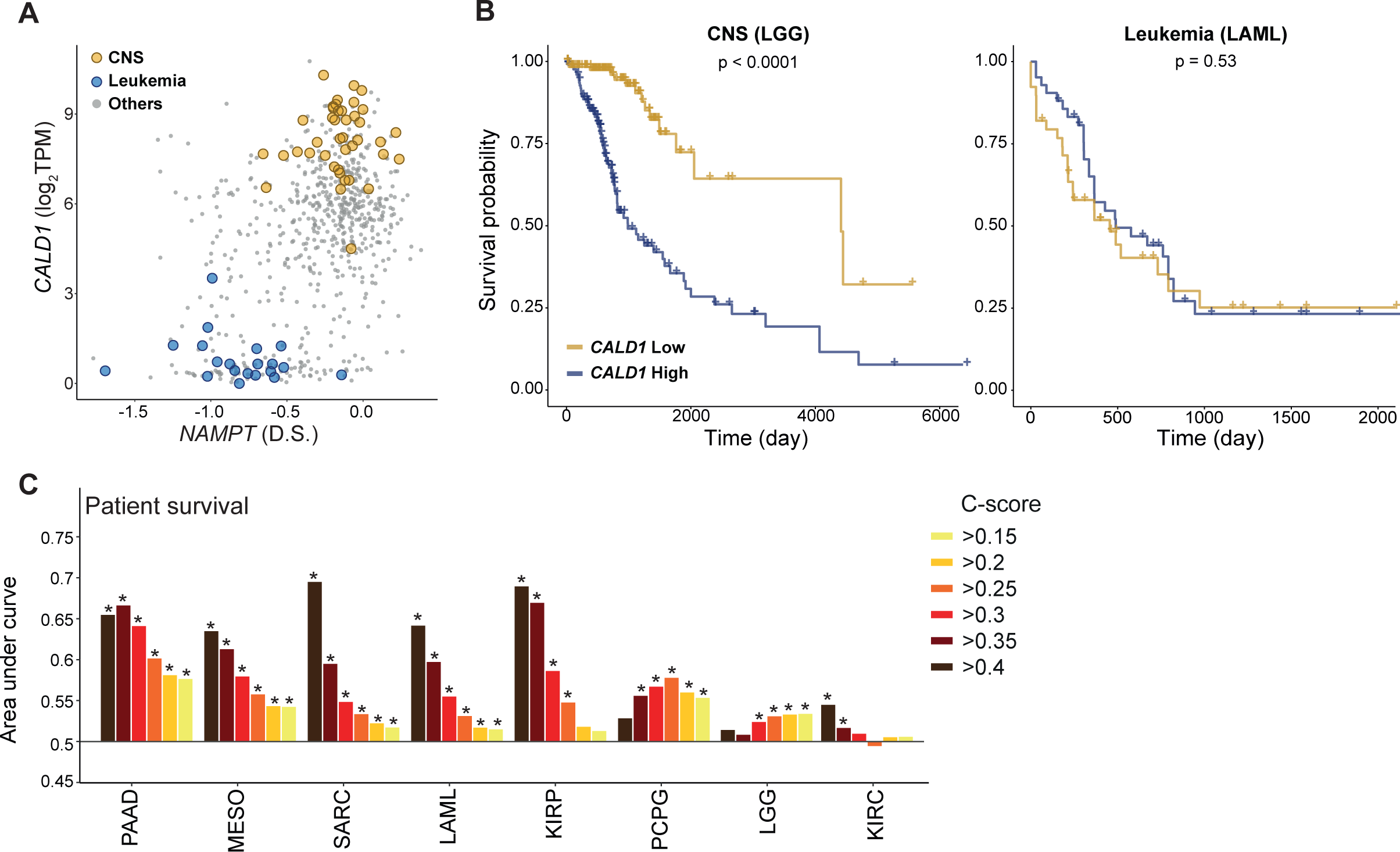
Harnessing cell-specific high C-score gene pairs for cancer patient prognosis. (A) C-score plot of *NAMPT* dependency score and *CALD1* gene expression (C-score = 0.447). Yellow circles represent central nervous system (CNS) cell lines and blue circles denote leukemia cell lines. (B) Kaplan-Meier overall survival plots for CNS (LGG, lower grade glioma, left panel) and leukemia (LAML, acute myeloid leukemia, right panel) cancer patients. Patients were stratified by high (>75%) or low (<25%) expression of *CALD1*, and *p*- values were calculated using Cox regression controlling for age, sex, pathological staging, clinical staging, and tumor grade, and corrected for multiple testing (false discovery rate < 0.2). (C) AUC of ROC curves based on C-score gene pair-based prediction of survival for each cancer type with different C-score cutoffs. Only the cancer types with at least one significantly positive C-score cutoff and those containing more than 1% of genes predicting patient survival with statistical significance are shown (* denotes *p* < 0.05).

We systematically assessed how buffering capacity from C-score-identified gene pairs could help predict cancer patient survival for all 30 TCGA cancer types. We found that for 15 of those cancers, at least 1% of genes across the genome can predict patient survival (with statistical significance assessed by Mann-Whitney U test), and for 8 of those 15 cancer types, the performance of CEBU-mediated buffering capacity for at least one C-score cutoff was significantly better than random (AUC > 0.5, false discovery rate < 0.2) (**Fig. 6C**). In addition, CEBU-mediated buffering capacity is also predictive of pathological stage (**Fig. S6A**), clinical stage (**Fig. S6B**), and tumor grade (**Fig. S6C**) for multiple cancer types. In general, buffering capacity-based predictions performed better for higher C-score cutoffs. Taken together, our results show that the CEBU-mediated buffering capacity derived from our C-score index can be indicative of cancer aggressiveness, as illustrated by patient survival, cancer pathological stage, clinical stage and tumor grade.

## Discussion

In multicellular organisms, different cells and tissues conduct various functions via specialized cellular structures and/or according to specific states (e.g., signaling and/or metabolic states) by regulating cell- and tissue-specific gene expression. Our study indicates that this cell- and tissue-specific gene expression not only contributes directly to tissue-specific functions, but also allows buffering for functional enhancement. This type of functional buffering, which we have termed cell-specific expression buffering (CEBU), can enhance housekeeping functions in specific tissues, thereby enabling tissue homeostasis. Furthermore, it appears to be especially prevalent in tissues of low regenerative capacity (e.g., bone and neuronal tissues) and it can promote tumor aggressiveness based on cancer patient survival. Although functional buffering has long been known as critical to biological robustness, the mechanisms underlying functional buffering remain largely unknown ^3^. CEBU that we illustrate in the present study represents a possible buffering mechanism in multicellular organisms that is critical for tissue homeostasis and cancer robustness.

One key feature of CEBU is the distinct patterns of expression and dependency (essentiality) between the buffered genes (G1s) compared to buffering genes (G2). In general, G1s tend to be broadly expressed with stronger dependency, whereas expression of G2s is more tissue-specific and less essential (**Fig. 4A and S2C**). Generally, the essentiality of genes is correlated with their expression level and tissue specificity ^31–33^. Housekeeping genes that are broadly expressed in most cells exhibit stronger essentiality. In contrast, genes expressed in specific cell types are considered to have weaker essentiality. Here, CEBU represents a putative mechanistic link between these two types of genes (i.e., housekeeping and tissue-specific genes), enabling their cooperation to regulate cellular functions via functional buffering. Specifically, house-keeping functions like metabolism, transcription, translation, and cell-cycle-related processes are highly enriched among high C-score gene pairs (**Fig. 2D**), indicating that house-keeping functions can be robustly maintained via CEBU-mediated functional buffering.

As a cell- and tissue-specific buffering mechanism, CEBU may endow buffering capacity on specific cells/tissues in order to maintain their functions and survival. This enhancement of cellular robustness may allow cells to persist for a longer time-period, in some cases even throughout the lifespan of an organism. As a result, CEBU may compensate for the lack of regenerative capacity in certain tissues. We predicted neuronal and bone tissues to have the strongest CEBU-mediated buffering capacities (**Fig. 4C** and **5C**), both of which exhibit relatively low regenerative capacities ^34–36^. In contrast, human blood cells, which are fully regenerated in 4 to 8 weeks ^37^, are predicted to have the weakest buffering capacities (**Fig. 5C**). Therefore, it is tempting to speculate that cell types of weaker regenerative capacities, such as neurons, need to sustain robust cellular functions through the buffering afforded by CEBU, thereby maintaining their tissue homeostasis. In contrast, highly regenerative tissues are frequently replaced, so they have less need for functional buffering.

Unlike the needs-based buffering mechanism, whereby the buffering gene is only activated when its buffered function is compromised, the CEBU-mediated intrinsic buffering proposed herein maintains a constitutively active state with cell- and tissue-specificity. Since the buffering gene (G2) is continuously expressed, there is no need for a control system to monitor if a function has been compromised and to activate the expression of the buffering genes. As a result, no response time is needed for intrinsic buffering, unlike for needs-based buffering. Accordingly, the CEBU mechanism can enable or adjust buffering capacity by regulating the expression of buffering genes via cell- or tissue-specific epigenetic regulators. Thus, CEBU can buffer housekeeping functions that need to be performed constitutively, which differs from the needs-based buffering that is mostly characterized as stress-responsive ^38^. Overall then, CEBU describes a simple, efficient and potentially versatile mechanism for functional buffering in humans and potentially other multicellular organisms.

CEBU describes an intrinsic buffering mechanism that functions under normal physiological conditions. Consistent with this notion, when we examined if our C-score index could be biased due to our usage of cancer cell lines, we found that only a low percentage (2.3% per gene pair, **Fig. S7A**) of mutant cell lines contributed to our C-score measurements. Moreover, excluding mutant cell lines did not qualitatively affect our C-score measurements, especially for high C-score gene pairs (**Fig. S7B**). The same trend holds for cancer-related genes (**Fig. S7B**). These results indicate that mutant cell lines are not the major determinants of C-scores. Similarly, since copy number variation (CNV) is a major mechanism for oncogenic expression, we checked if CNV contributes to G2 expression. As shown in **Figure S7C**, the correlation between G2 expression and copy number decreases with increasing C-score, indicating that CNV is not a primary mechanism regulating G2 expression. Thus, our C-score index is likely not biased by the utilization of cancer cell lines.

We observed an enrichment of duplicated genes among high C-score gene pairs, supporting the notion that duplicated genes contribute to the context-dependent essentiality of their paralogous genes ^11–13^. In addition to duplicated genes, our C-score index identified a high percentage of non-duplicated gene pairs with high buffering capacities (**Fig. S3A**), and these non-duplicated gene pairs tend to belong to the same pathways and/or protein complexes (**Fig. 2B** and **2C**). Therefore, it is possible that many of these G1s and G2s represent non-orthologous functional analogs. One simple scenario could be that G1 and G2 physically interact with each other to form a protein complex, wherein G1’s function can be structurally substituted by G2. Indeed, we identified the *POP7* and *RPP25* gene pair as an example of this scenario (**Fig. 3C** and **3D**). More sophisticated and indirect functional buffering can also occur between G1s and G2s given the complex interactions among biological functions ^39^.

We expect that CEBU exerts buffering effects through additional types of molecular interactions, which remain to be tested experimentally.

C-score-derived cell-specific buffering capacities comply well with experimentally validated genetic interactions in human cells (**Fig. 5B** and **Fig. S5**), indicating that CEBU may represent a critical mechanism for synthetic lethality in human cells. In practice, it remains a daunting challenge to systematically characterize genetic interactions in organisms with complex genomes due to large numbers of possible gene pairs, i.e. ∼200 million gene pairs in humans. Previous efforts have used computational approaches on conserved synthetically lethal gene pairs in the budding yeast *Saccharomyces cerevisiae* or employ data mining on multiple large datasets to infer human synthetic lethalities ^40, 41^. Nevertheless, predictions emanating from different studies exhibit little overlap ^42^, evidencing the marked complexity of synthetic lethality in humans. The CEBU mechanism proposed here can contribute both experimentally and computationally to a better characterization of human genetic interactions.

Using G2 expression of a high C-score gene pair to stratify cancer patients, we observed a significant difference in cancer patient survival, indicating that stronger CEBU-mediated buffering capacity could be predictive of cancer aggressiveness in patients (see **Fig. 6A** and **6B** for an example). Indeed, buffering capacity helped predict cancer patient survival in 8 of 15 cancer types and, generally, predictive performance was better for higher C-score cutoffs (**Fig. 6C**). Apart from patient survival, buffering capacity is also indicative of pathological stage, clinical stage, and tumor grade of cancers (**Fig. S6**). These results support our hypothesis that stronger buffering capacity via higher G2 expression contributes to cancer robustness in terms of proliferation and drug resistance. Given the complexity of cancers, it is surprising to see such general predictivity of cancer prognosis by individual high C-score gene pairs. Accordingly, we suspect that some cancer cells may adopt this cell- and tissue-specific buffering mechanism to enhance their robustness in proliferation and stress responses by targeting the expression of buffering genes. Clinically, the expression of such buffering genes could represent a unique feature for evaluating cancer progression when applied alongside other currently used clinical characteristics. Finally, experimental validation of C-score-predicted genetic interactions will help identify potential drug targets for tailored combination therapy against specific cancers.

## Materials and Methods

### Retrieval and processing of dependency score and gene expression data

Data on dependency scores and CCLE (Cancer Cell Line Encyclopedia) gene expression were downloaded from the DepMap database (DepMap Public 19Q4) ^15, 16^. Dependency scores modeled from the CERES computational pipeline based on a genome-wide CRISPR loss-of-function screening were selected. CCLE expression data was quantified as log_2_ TPM (Transcripts Per Million) using RSEM (RNA-seq by Expectation Maximization) with a pseudo-count of 1 in the GTEx pipeline (https://gtexportal.org/home/documentationPage). Only uniquely mapped reads in the RNA-seq data were used in the GTEx pipeline. Integrating and cross-referencing of the dependency score and gene expression datasets yielded 18239 genes and 684 cell lines. Genes lacking dependency scores for any one of the 684 cell lines were discarded from our analyses.

### C-score calculation

Our C-score index integrates the dependency scores of buffered genes (G1) and the gene expression of buffering genes (G2) to determine the buffering relationship between gene pairs. Genes with mean dependency scores > 0 or mean gene expression < 0.5 log_2_ TPM were discarded, yielding 9196 G1s and 13577 G2s. The C-score integrates the correlation (*ρ*) and slope between the dependency score of gene *G*1 and the gene expression of gene *G*2, defined as:

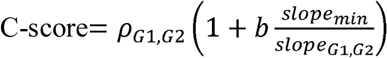

where *ρ* denotes the Pearson correlation coefficient and *slope_min_* denotes the minimum slope of all considered gene pairs that present a statistically significant positive correlation. The normalized slope can be weighted by cell- and tissue-type specific *b*. In our analysis, *b* is set as 1 for a pan-cell or pan-cancer analysis.

### Duplicated gene assignment

Information on gene identity was obtained from ENSEMBL (release 98, reference genome GRCh38.p13) ^43^. Two genes are considered duplicated genes if they have diverged from the same duplication event.

### Enrichment analysis for buffering gene pairs

For enrichment analysis of gene pairs, we adopted a previously described methodology ^44^. Briefly, GO and KEGG gene sets were downloaded from the Molecular Signatures Database (https://www.gsea-msigdb.org/gsea/msigdb/). The number of total possible gene pairs is 9196 (G1) x 13577 (G2). The condition of G1 and G2 being the same gene was excluded as a potential buffering gene pair under all C-score cutoffs. Enrichment was calculated as:

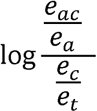

Where represents the number of gene pairs that are both annotated and with buffering capability, is the number of annotated gene pairs, is the number of buffering gene pairs, and is the total number of gene pairs.

Protein-protein interaction (PPI) data was downloaded from the STRING database (version 11) ^20^. Only high-confidence interactions (confidence > 0.7) in human were considered. The STRING database determines confidence by approximating the probability that a link exists between two enzymes in the KEGG database. Data on protein core complexes were downloaded from CORUM (http://mips.helmholtz-muenchen.de/corum). The enrichment calculation is the same as for GO and KEGG, except that represents the number of gene pairs that have PPI or are in the same complex and have buffering capability, and is the number of gene pairs that have PPI or are in the same complex.

### Construction of our Human Compensatory Gene Network

The directional human compensatory gene network was constructed from gene pairs exhibiting high C-scores (> 0.25). For illustration, isolated subnetworks are not shown. We visualized the network using Cytoscape (https://cytoscape.org/) and MATLAB. GO enrichment was conducted on each cluster using g:Profiler ^45^. To identify functionally-related gene clusters in the human compensatory gene network, the genes with enriched functions were inputted into the SAFE algorithm ^46^. The neighbor radius was determined by regional enrichment of sub-networks for each GO-enriched function.

### Experimental validation

A549, H4, HT29, LN18, MCF7, and U2OS cell lines were selected based on their distribution across the C-score plots (**Fig. 3A and 3** ), indicating different buffering capacities. All cell lines were purchased from ATCC and they were cultured in Dulbecco’s Modified Eagle Media (H4 and LN18), Ham’s F-12K Medium (A549), or RPMI 1640 media (HT29, MCF7, and U2OS) supplemented with 5% fetal bovine, serum, 100 U/mL penicillin, 100 μg/mL streptomycin, and 250 ng/mL fungizone (Gemini Bio-Products). Cell growth was monitored by time-lapse imaging using Incucyte Zoom, taking images every 2 hours for 2-4 days. To suppress *FAM50A, FAM50B*, *POP7* and *RPP25* expression, lentivirus-based shRNAs were delivered individually or in combination. The gene-specific shRNA sequences are: *FAM50A* - CCAACATTGACAAGAAGTTCT and GAGCTGGTACGAGAAGAACAA; *FAM50B* – CACCTTCTACGACTTCATCAT; *POP7* – CTTCAGGGTCACACCCAAGTA and CGGAGACCCAATGACATTTAT; and *RPP25* – CCAGCGTCCAAGAGGAGCCTA.

To ensure better knockdown of gene expression, shRNAs were delivered twice (7 days and 4 days before seeding). Equal numbers of cells were seeded for cell growth measurements by time-lapse imaging using Incucyte Zoom. The lentivirus-based shRNAs were purchased from the RNAi core of Academia Sinica. The growth rate under each condition was measured by fitting cell confluence to an exponential growth curve using the Curve Fitting Toolbox in MATLAB.

### Bliss independence model

Cytotoxic synergy was measured using the Bliss independent model ^22^. The Bliss model is presented as a ratio of the expected additive effect to the observed combinatorial effect:

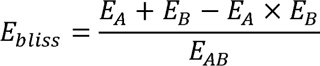

where *E* is the effect of drug *A*, *B*, or a combination of *A* and *B*. Effect was measured by the relative cell growth, based on the fold-change of confluency between 0 and 72 hours upon suppression of *FAM50A* and *FAM50B* or suppression of *POP7* and *RPP25* in all cell lines except MCF7, and between 0 and 96 hours upon suppression of *FAM50A* and *FAM50B* in MCF7.

### Cell-specific buffering capacity and comparison to experimental genetic interactions

Cell-specific buffering capacity was derived from the C-score of a given gene pair and gene expression of the buffering gene (G2) in the cell line of interest following the equation:

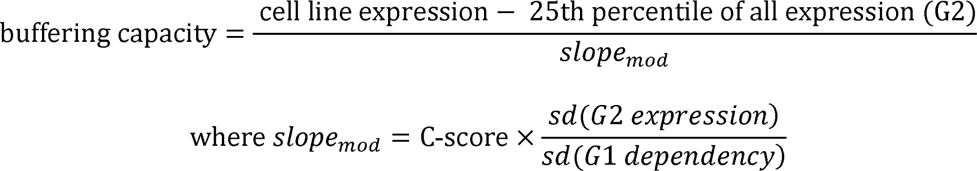

where *sd* = standard deviation. The 25^th^ percentile cutoff for expression is determined empirically, although different percentile cutoffs do not qualitatively affect the measurements of buffering capacities (**Fig. 5A**).

Combinatorial CRISPR screen-derived genetic interaction scores were pooled from four literature sources ^25–28^ (**Table S1**). We only considered cell lines that appear in DepMap CERES 19Q4. There were two C-scores for each gene-pair of the experimental dataset (either gene could be a G1), and we assigned the higher C-score for that gene-pair. Overall, we curated 10,222 genetic interaction scores in various cell lines from the literature, and 1986 out of 10,222 genetic interaction scores had a C-score > 0.1. To evaluate the validity of buffering capacity, we generated a ground-truth dataset by assigning gene-pairs with a positive genetic interaction as false for buffering and a negative genetic interaction as true for buffering. The qualitative performance of buffering capacity against this ground-truth dataset was assessed by ROC curve. Additionally, we correlated the buffering capacity directly via a ground-truth genetic interaction score for quantitative evaluation.

### Tissue specificity

To calculate tissue-specificity, cell lines were grouped by their respective tissues, and expression of genes in cell lines of the same tissue were averaged. Tissue specificity was calculated as tau (*τ*) ^24^, where *τ* is defined as:

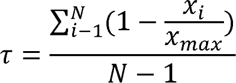

with *N* denoting the number of tissues, *x_i_* denoting the expression of a gene, and *x_max_* denoting the highest gene expression across all tissues. Note, expression values were log-transformed, so log_2_ TPM < 1 was considered as 0 in tissue specificity calculations ^47^.

### Cancer-specific survival prediction according to C-score gene pairs

Gene expression and survival data from The Cancer Genome Atlas (TCGA) ^29^ was retrieved from Xena ^48^. The DepMap cancer cell lines were mapped to TCGA cancers based on the annotation in **Table S2** (cancers that do not have a matched cancer type in CERES 19Q4 were not analyzed). To systematically analyze cancer prognosis, we first performed a multiple test correction on the *p*-values from Cox regression controlling for age, sex, pathological stage, clinical stage and tumor grade. We calculated the false discovery rate (FDR) using the Benjamini–Hochberg procedure with a threshold < 0.2. The ground-truth table for each cancer was constructed using the adjusted *p*-value. AUC of ROC curves were used to assess the performance of survival based on buffering capacity. AUCs and ROCs were generated using python and R. The statistical significance of AUC was assessed by Mann-Whitney U test ^49^ to evaluate if gene expression with a positive Cox coefficient (poorer prognosis) reflected significantly higher buffering capacities in each cancer with different C-score cut-offs. The *p*-values of the Mann-Whitney U test were adjusted using the Benjamini-Hochberg procedure with a threshold < 0.2. We conducted a similar approach to the prognosis analysis for buffering capacities and clinical features. We calculated the p-values of correlations between gene expression and clinical features, staging and grade, and corrected the p-values using the Benjamini-Hochberg procedure with a threshold < 0.2. We then used Mann-Whitney U tests to evaluate if gene pairs with a significant positive correlation between gene expression and tumor aggressiveness presented a significantly higher buffering capacity in each cancer for different C-score cut-offs.

### Data Availability

All high C-score (> 0.25) gene pairs (https://figshare.com/s/6f8929c6543687a6062f) and programming code (https://figshare.com/s/b778489bb2f6fc3b0069) are available in the FigShare repository.

## Acknowledgement

We thank members of the Lab for Cell Dynamics for helpful discussions. We are grateful to Jose Reyes, Hannah Katrina Co, and Jun-Yi Leu for their comments and suggestions on the manuscript, and Ann Mikaela Lynne Ong Co, Su-Ping Lee, the Imaging Core at the Institute of Molecular Biology, and the RNAi Core at Academia Sinica for their technical support.

## Author Contributions

H.-K.L., J.-H.C., and S.-h.C. conceived the project. H.-K.L., J.-H.C., C.-C.W., and S.-h.C. designed and conducted the computational and statistical analyses. H.-K.L., J.-H.C., C.-C.W., and S.-h.C. wrote the manuscript. F.-S.H., C.D. and S.-h.C. designed the experiments, and F.- S.H., and C.D. conducted the experiments.

## Supporting Information Captions

Figure S1. Distribution of C-scores

Figure S2. Correlation properties of G1 dependency score and G2 gene expression according to increasing C-score

Figure S3. Functional and pathway enrichments of C-score-identified duplicated genes

Figure S4. Shuffling the G2-tissue relationship disrupts G2 tissue-specificity

Figure S5. C-score-based prediction of cell-specific genetic interaction using buffering capacity

Figure S6. C-score-based prediction of cancer pathological stage, clinical stage and tumor grade

Figure S7. Decreasing effects of mutational variation as C-score increases

Table S1. List of combinatorial CRISPR-screened gene pairs in published literature

Table S2. Cross-reference table for TCGA cancer type and DepMap cancer type

