## Supplementary Methods and Figures for "Functional buffering via cell-specific gene expression promotes tissue homeostasis and cancer robustness"

Supplementary Material

**Supplementary Methods**

**Construction of C-score null distribution and modeling as a normal distribution**

As a control, expression of each gene was randomly shuffled amongst the 684 cell lines. The shuffled expression values were used in place of true expression in the C-score calculation to create a randomly shuffled null distribution. Five shuffled expression datasets were generated to calculate five null distributions, one of which is shown. These null distributions fit a normal distribution based on qqplot analysis. The mean and standard deviation of the null distributions were used to model a normal distribution and statistical significance was assessed using a *t*-test.

**Control for mean and standard deviation for dependency and expression analysis**

To analyze the increase in standard deviation of dependency/expression with increasing C-scores, we controlled for the mean value of dependency/expression. For each gene, 100 other genes with mean dependency/expression differing the least from that gene were selected from the genome. Then, one gene was randomly selected from out of those 100 genes to be the control for mean. The distribution of the standard deviations of the controls was plotted alongside the original distribution. This process was repeated five times to ensure that there was no selection bias during random selection. Only one of the five replicates is shown.

**Mutational analysis**

Mutation information was retrieved from DepMap. We considered all types of mutations, including hotspot, damaging, and non-conserving mutations. Genes with significant oncopotential^1^ to be oncogenes or tumor suppressors were considered cancer-related genes, otherwise they were considered non-cancer-related.

**Copy number analysis**

Copy number was retrieved from DepMap. We used bootstrapping to estimate the random correlation of copy number variation and expression while controlling for expression mean and standard deviation. Genes were divided into 100 bins based on their expression mean and standard deviation, respectively. Each G2 was randomly replaced by a gene in the same bin. We performed 10,000 randomizations under each C-score cutoff for every G2.

**Supplementary Figure and Figure Legends**

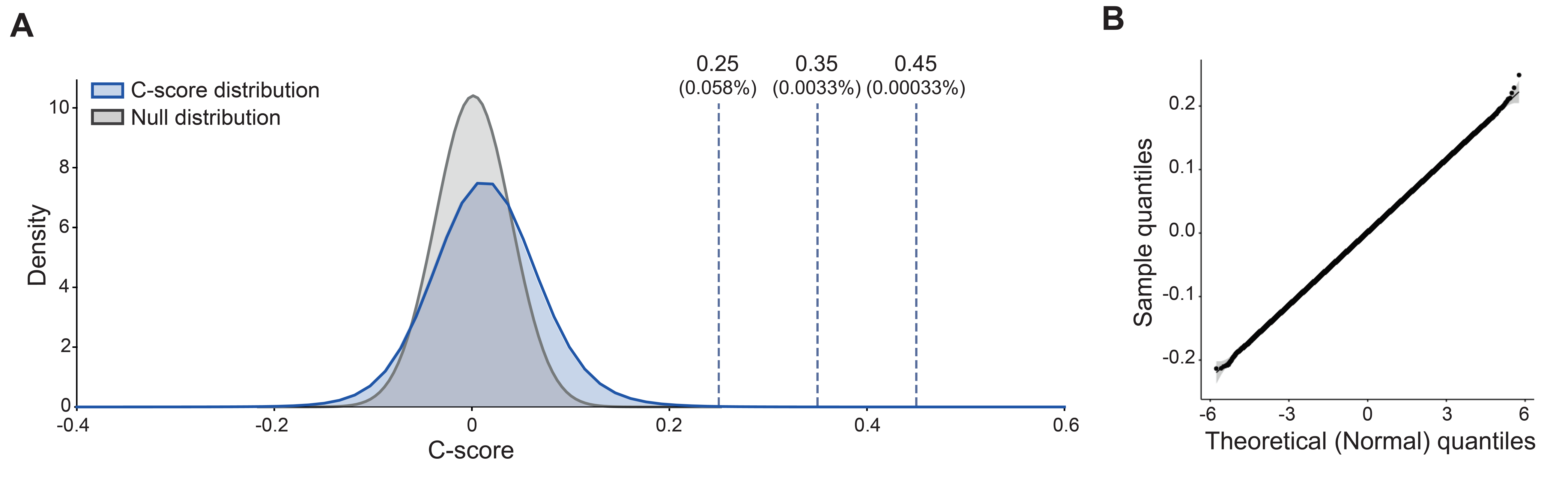

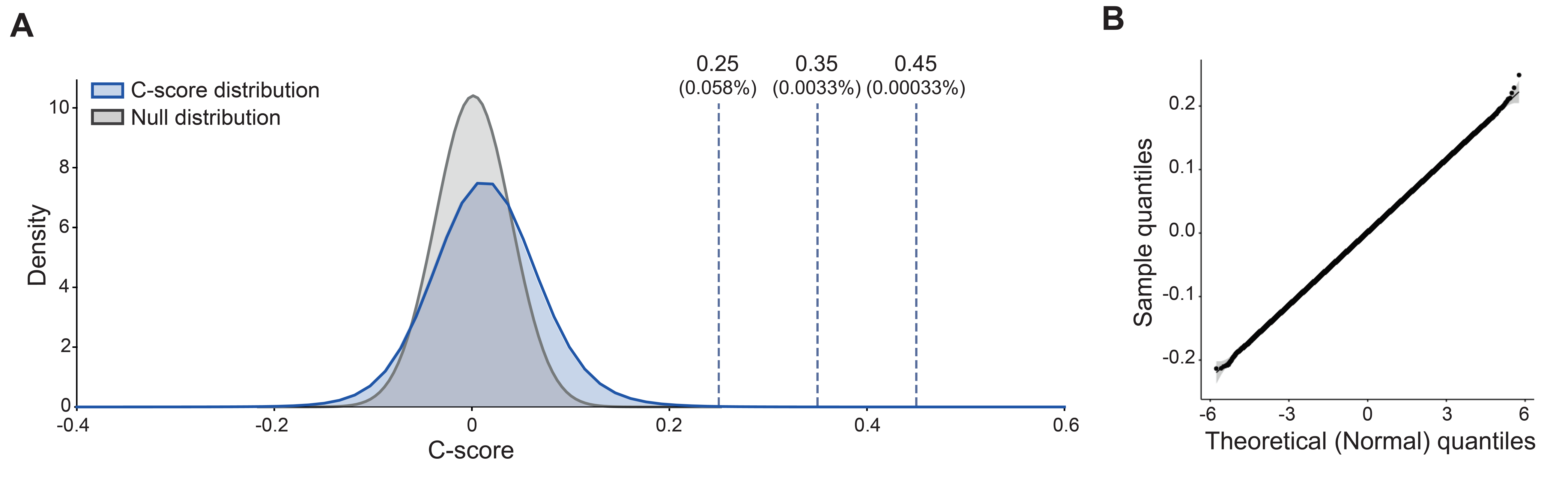

**Supplementary Figure 1. Distribution of C-scores**

(A) Distribution of C-scores for all gene pairs (blue) and one of five randomly shuffled distributions (grey). The plot shows C-score values of between -0.4 and 0.6. The dashed lines indicate the C-score percentiles of 0.25, 0.35 and 0.45 for the all-gene-pair distribution. (B) The randomly shuffled distribution represents a normal distribution generated using qqplot with a theoretical normal distribution. One of five qqplot distributions is shown. Range of mean and standard deviation for the five randomly shuffled distributions are 0.001-0.000 and 0.0383-0.0389, respectively.

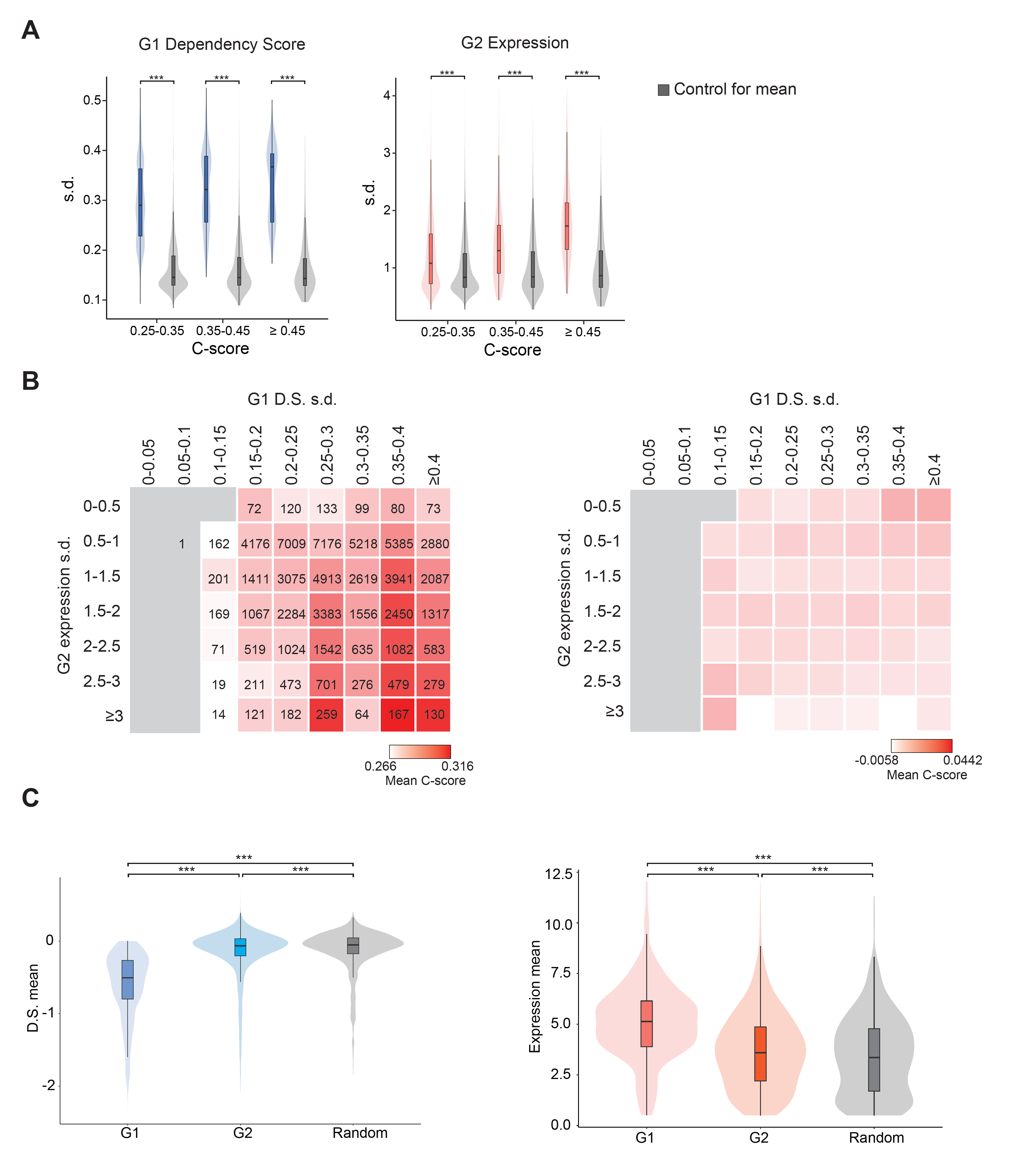

**Supplementary Figure 2. Correlation properties of G1 dependency score and G2 gene expression according to increasing C-score**

(A) The standard deviation (s.d.) of G1 dependency score (D.S.) and G2 gene expression increases with increasing C-score. Grey represents control distributions for mean values, which were constructed based on the s.d. of randomly selected genes with similar mean values of dependency or expression (see **Supplementary Methods**). Differences between each C-score interval to the control distribution for mean values are significant based on a one-tailed two-sample *t*-test (*** denotes *p*-value < 0.001). Five random selections were performed, and the results are comparable. One of the five random selections is shown. (B) Left panel: mean C-score of intervals of increasing s.d. for G1 dependency and for G2 expression values. Numbers of genes in each interval are shown in the cells. Only cells containing more than 10 genes are colored. Right panel: mean C-score of intervals of increasing s.d. of G1 dependency and randomly shuffled G2 expression values. Random shuffling of G2 expression ensured the same mean and s.d. values were retained, but it uncoupled the cell line associations, which acted as a control for the increased s.d. of both G1 dependency and G2 expression with increasing C-score. (C) Mean of dependency score (D.S., blue) and gene expression (red) of G1 and G2. The random plot (grey) was constructed by randomly selecting the same number of genes from the genome (*** statistically significant at *p* < 0.001, based on a one-tailed paired *t*-test for G1 and G2 comparison and one-tailed two-sample *t*-test for comparison with random).

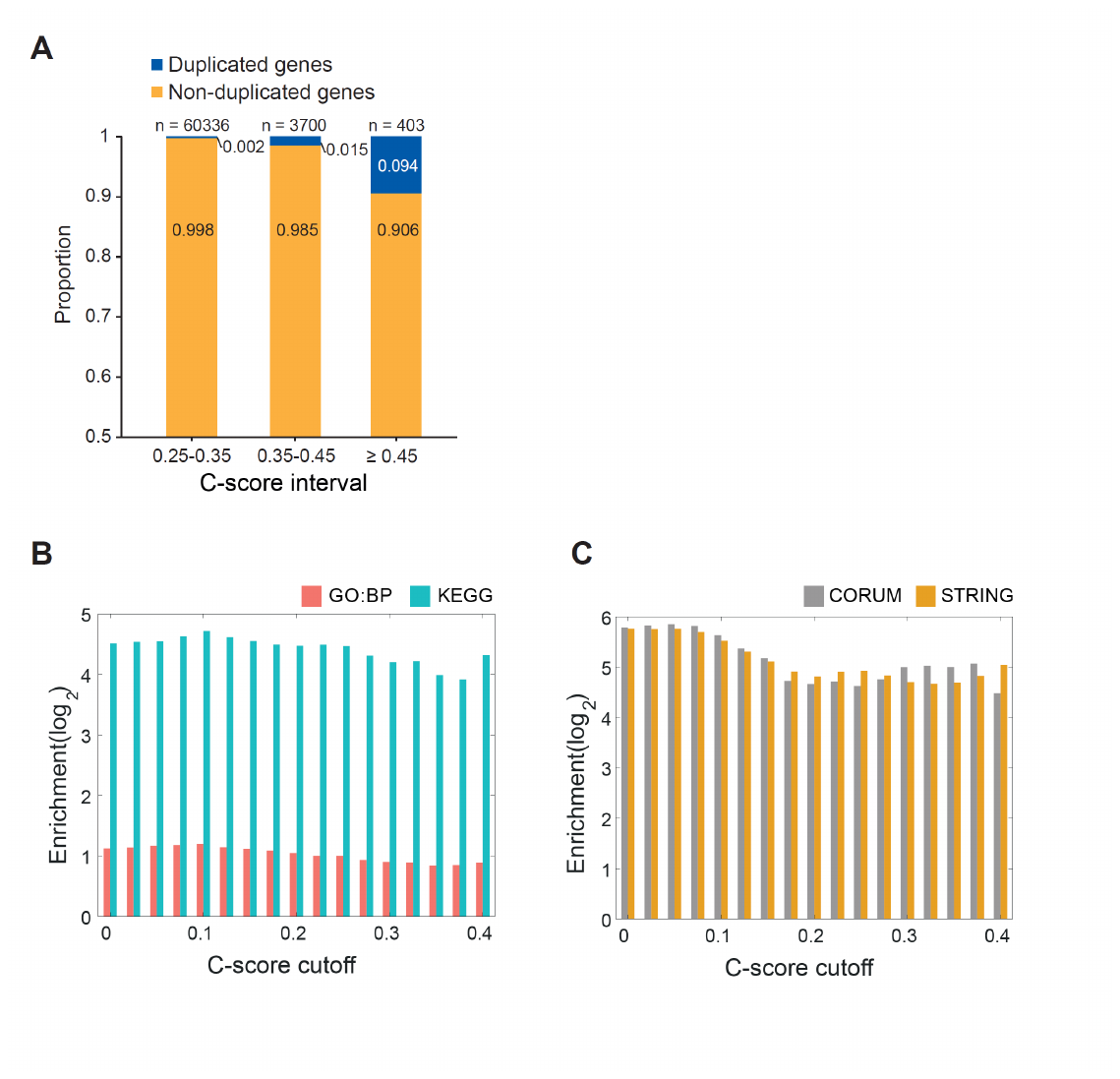

**Supplementary Figure 3. Functional and pathway enrichments of C-score-identified duplicated genes**

(A) Percentage of C-score-identified non-duplicated and duplicated gene pairs with different C-score cut-offs. (B-C) Functional enrichment of C-score gene pairs increases with C-score cutoff for duplicated genes. (B) Pairs of duplicated genes annotated with the same gene ontology biological process (GO:BP) term or KEGG pathway are enriched in C-score gene pairs, and enrichment remains relatively constant with increasing C-score cut-off. (C) Pairs of duplicated genes with annotated protein-protein interactions from STRING and within the same protein complex from CORUM are enriched in C-score gene pairs, and enrichment remains relatively constant with increasing C-score cut-off.

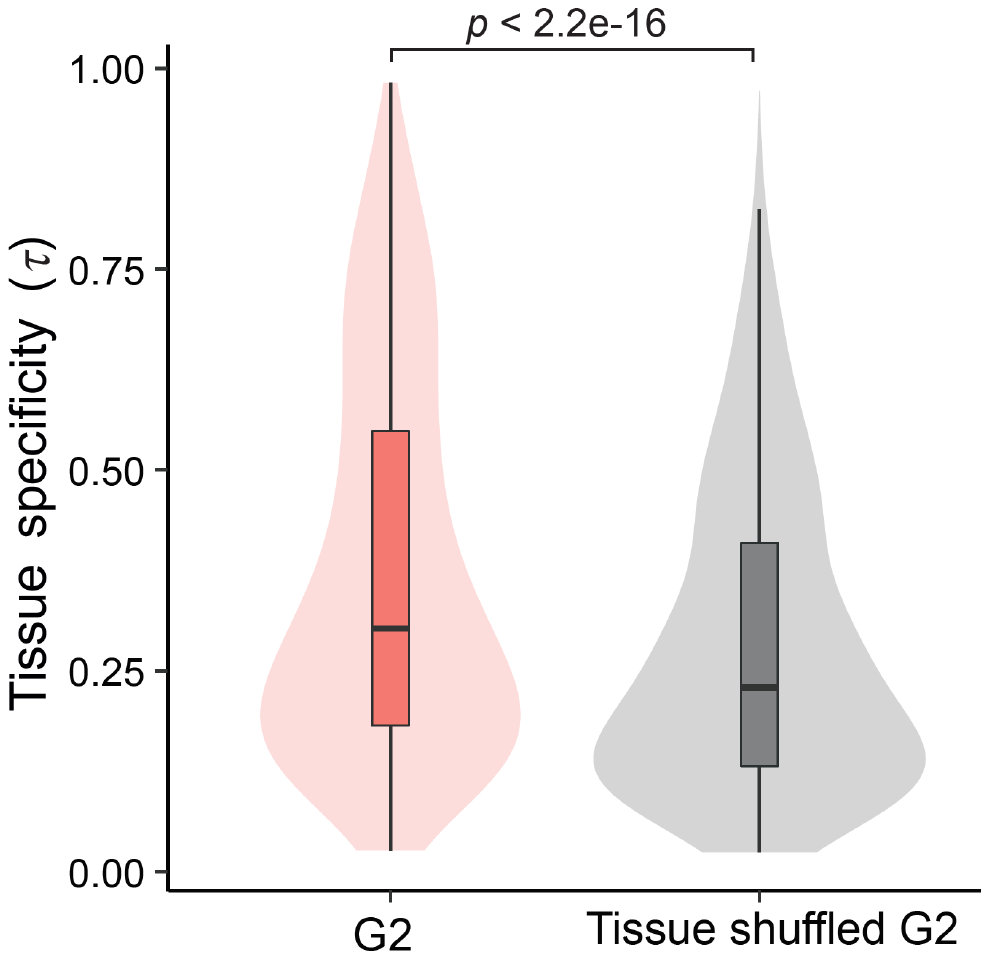

**Supplementary Figure 4. Shuffling the G2-tissue relationship disrupts G2 tissue-specificity**

Tissue specificity (*τ*) of G2s and after we had randomly shuffled the tissues for the cell lines associated with G2s. *τ* was calculated for tissue-shuffled G2s from high C-score gene pairs. Statistical significance was assessed by paired-*t­* test.

**Supplementary Figure 5. C-score-based prediction of cell-specific genetic interaction using buffering capacity**

(A) Correlation (*ρ* = −0.231, *p* = 0.034, n = 84) between existing published genetic interactions (see **Table S1** for detailed information) and our predicted buffering capacities with a C-score cutoff of 0.25. (B) Concordance (*ρ* = −0.945, *p* < 0.05) between different C-score cutoffs and the correlation (shown in A) between existing published genetic interactions and our predicted buffering capacities. (C) Effect of varying the G2 expression percentile cutoff (from 20% to 50%) on the correlations between our buffering capacities and experimental results according to different C-score cut-offs.

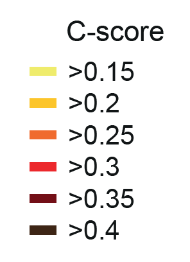

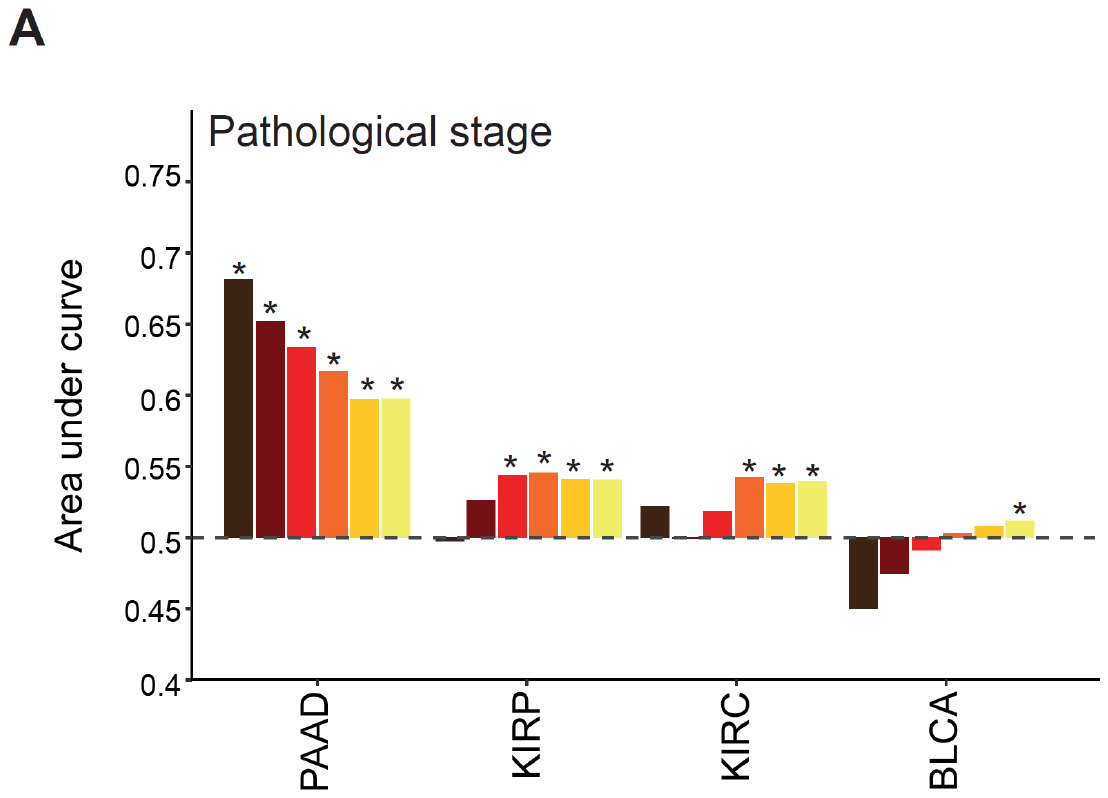

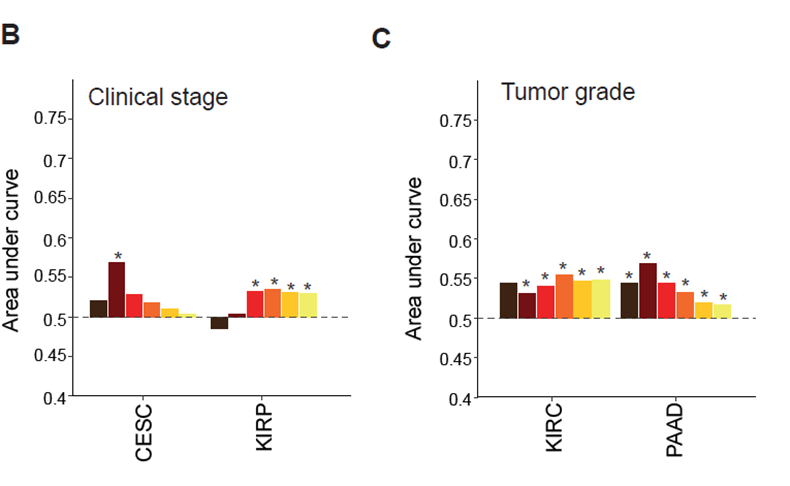

**Supplementary Figure 6. C-score-based prediction of cancer pathological stage, clinical stage and tumor grade**

Area under a curve (AUC) of ROC curves illustrating predictive performance for (A) pathological stage, (B) clinical stage, and (C) tumor grade after multiple testing correction (false discovery rate < 0.2) using different C-score cutoffs (>0.15 to >0.4). Only AUCs > 0.5, the cancer types with at least one significantly positive C-score cutoff, and cancers containing more than 1% of genes predicting patient survival with statistical significance are shown. *denotes false discovery rate < 0.2.

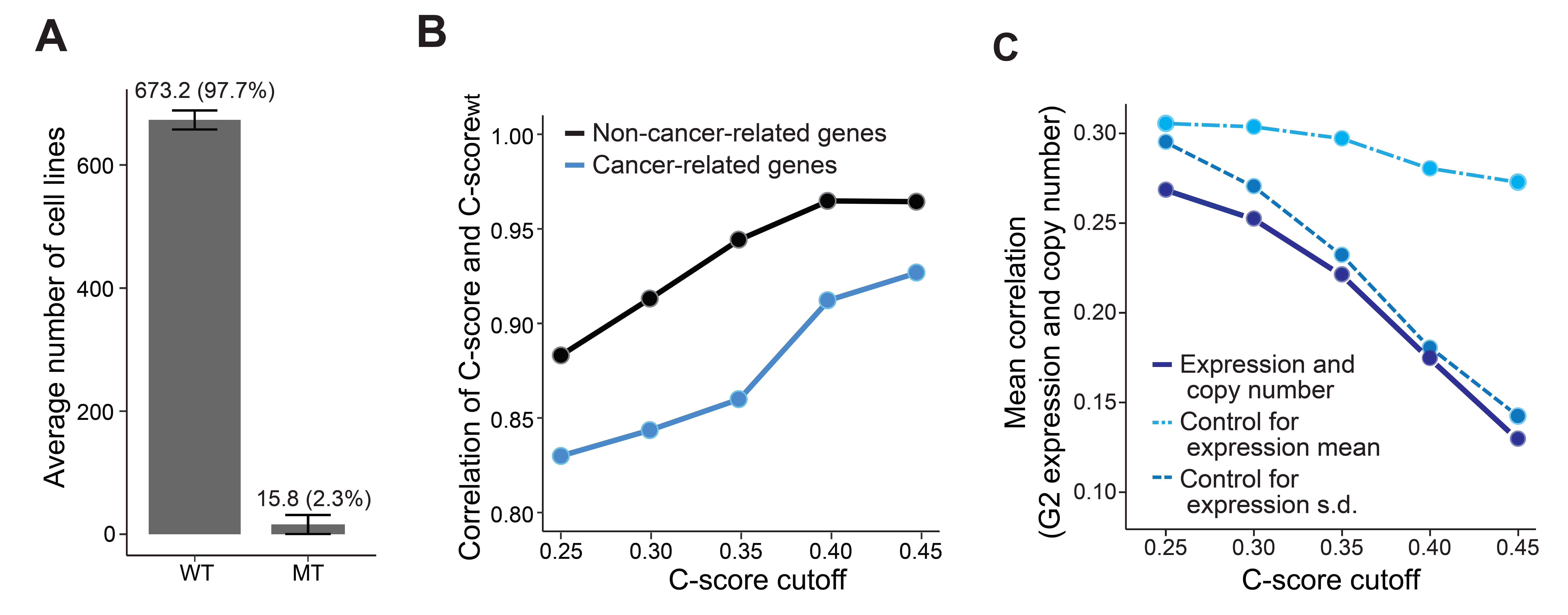

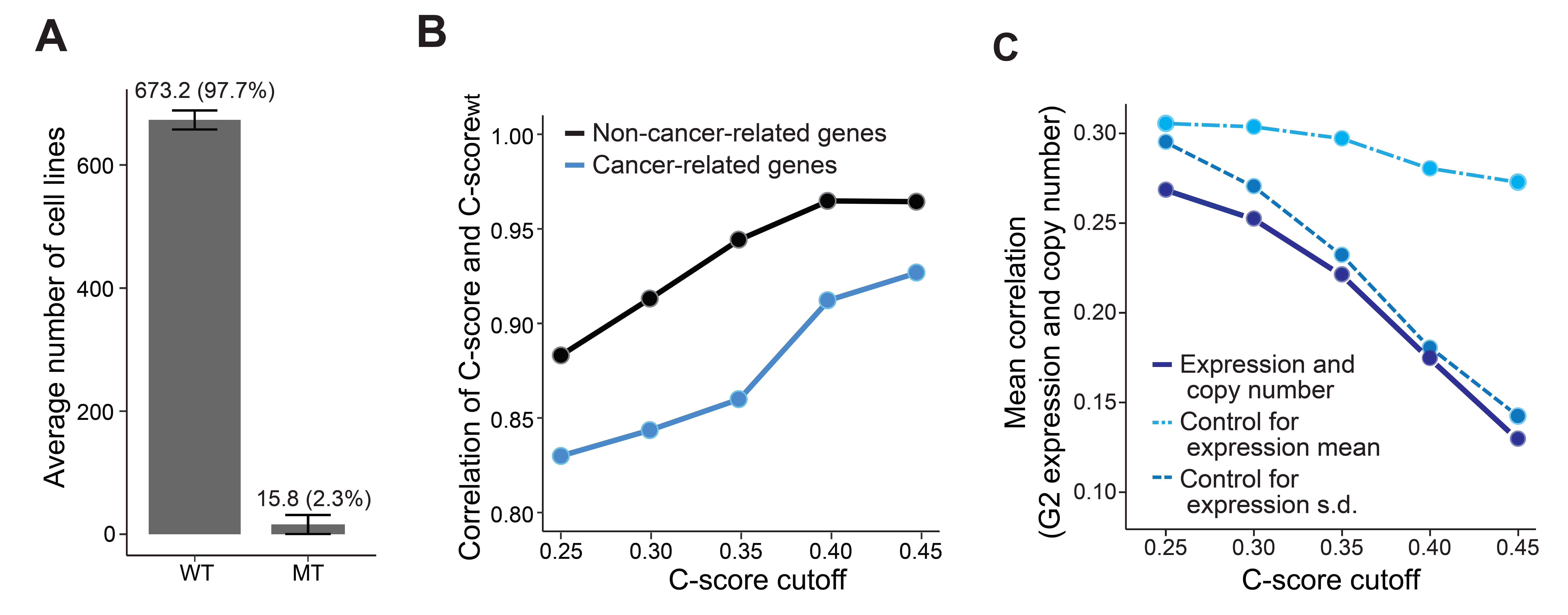

**Figure S7. Decreasing effects of mutational variation as C-score increases**

(A) Average number of wild type or mutated cell lines for each C-score gene pair. (B) Correlation between C-scores and C-scores calculated using only wild-type genes (C-score_wt_) for different C-score cutoffs. Genes were assigned as cancer-related if they have a significant oncopotential^1^ to be oncogenes or tumor suppressor genes. Black line represents non-cancer-related genes. Blue line represents cancer-related genes. (C) The mean correlation between G2 copy number and its expression for different C-score cut-offs is represented by the dark blue line. The lighter blue dashed lines are correlation controls for G2 expression mean and standard deviation. Controlling for G2 expression mean and standard deviation revealed that standard deviation is the major contributor to the diminished correlation between G2 expression and CNV.
